# Co-designed land-use scenarios and their implications for storm runoff and streamflow in New England

**DOI:** 10.1101/847186

**Authors:** Andrew J. Guswa, Brian Hall, Chingwen Cheng, Jonathan R. Thompson

## Abstract

Future changes in both landscape and climate have the potential to create or exacerbate problems with stormwater management, high flows, and flooding. In New England, four plausible land-use scenarios were co-developed with stakeholders to give insight to the effects on ecosystem services of different trajectories of socio-economic connectedness and natural resource innovation. To assess the effects of these land-use scenarios on water-related ecosystem services, we applied the Soil and Water Assessment Tool to two watersheds under two climates. Differences in land use had minimal effects on the overall water balance but did affect high flows and the relative contribution of storm runoff to streamflow. For most of the scenarios, the effect was small and less than the effect due to climate change. For one scenario – envisioned to have global socio-economic connectedness and low levels of natural-resource innovation – the effects of land-use changes were comparable to the effects due to climate. For that scenario, changes to the landscape increased the annual maximum daily flow by 10%, similar to the 5-15% increase attributable to climate change. These results, which were consistent across both watersheds, can help inform planning and policies regarding land use, development, and maintenance of hydrologic ecosystem services.

**Research highlights:**

1. Stakeholder-engaged scenarios provide meaningful and plausible futures for the New England landscape and assessment of effects of land-use change on storm runoff and streamflow
2. Effects of land use on the overall water balance are small across the landscape scenarios
3. Future land-use change has the potential to affect storm runoff and high flows to a degree that is comparable to the effects due to changes in climate in 2060
4. The degree of natural resource innovation affects storm runoff and high flows when population growth is large and has a negligible effect when population growth is low

## 1 Introduction

Changes to the landscape will affect water-related ecosystem services, and planning and development must be informed by the range of potential effects to ensure resilient and sustainable water resources, especially under a changing climate. While climate and precipitation and are the primary drivers of the hydrologic cycle, land use and land cover modulate those signals and can exacerbate or mitigate the impacts (Brauman et al., 2007). Watersheds concentrate precipitation inputs in space (regulating service), distribute them in time (regulating service), and remove water via evapotranspiration (provisioning service). Through modifications to infiltration capacity and vegetation cover, changes to land use and land cover will affect the partitioning of water between evapotranspiration and streamflow along with the timing of streamflows. An understanding of how plausible future landscapes and associated ecosystem services might affect the water balance and streamflow can improve planning, infrastructure design, and policy decisions.

An increase in vegetation cover tends to increase both evapotranspiration, which reduces the provision of streamflow, and infiltration, which increases the temporal regulation of streamflow. Paired watershed and observational studies generally show that a reduction in vegetation cover leads to an increase in average streamflow due to the reduction in evapotranspiration (e.g., Andréassin, 2004; Bosch & Hewlett, 1982; Brown et al., 2005; Brown et al., 2013; Bruijnzeel, 2004). In contrast, the effects of vegetation cover on low flows are less certain due to the competing effects on evapotranspiration and infiltration (e.g., Devito et al., 2005; Guswa et al., 2017; Homa et al., 2013; Jencso & McGlynn, 2011; Laaha et al., 2013; Price, 2011; Smakhtin, 2001). The direction of the effect on flooding and high flows is more certain, as vegetation increases evapotranspiration (reducing streamflow) and infiltration (reducing peak flows). The loss of vegetation, coupled with increases in impervious cover, increases peak flows. The magnitude and significance of those services are uncertain across environments and events, however. For example, in the UK, increases in vegetation were found to reduce peak flows for small to moderate rainfall events but had little effect for larger events (Dadson et al., 2017). When the land is saturated, the regulating effect of infiltration may be reduced, and some claim that landscape effects on flood reduction may be overestimated (Calder & Aylward, 2006). In one case, authors even found the opposite effect, with increased impervious area correlated with decreased high flows, perhaps due to a concomitant increase in stormwater detention infrastructure (Homa et al., 2013).

Investigators have also used modeling studies to elucidate the effects of land use on hydrologic ecosystem services. Karlsson et al. (2016) examined the combined effect of four land-use scenarios, four climate models, and three hydrological models on streamflows in Denmark and found that the climate model had more influence than land-use change. Ashage et al. (2018) used the Soil and Water Assessment Tool (SWAT) to show that forests and woodlands, relative to agriculture, regulated both sediment loads and peak flows in Tanzania. Baker & Miller (2013) also used SWAT in East Africa and found that increases in urbanization resulted in greater surface runoff and reduced groundwater recharge. For the Songkhram River Basin in Thailand, Shrestha et al. (2018) employed SWAT to determine that the effects of climate change (20% decrease in streamflow) were greater than the effects due to potential land-use changes (5% increase in streamflow). SWAT has also been applied to multiple watersheds in the United States. In the northeast, an increase in forest cover led to a decrease in the severity and duration of both high and low flows (Ahn & Merwade, 2017). In southern Alabama, Wang et al. (2014) showed that a near doubling of urban area from 26.4% of the landscape to 50.2% resulted in an increase of only 2.2% in the average daily flow. Hantush & Kalin (2006) simulated urbanization in the Pocono Creek in Pennsylvania, and they found that increasing development from 5.8% of the landscape to 75.8% reduced average flows by 1.1% and increased the average annual maximum daily flow by 19.4%. Cheng (2013) used SWAT to simulate and compare four land-use scenarios and three climate scenarios with respect to streamflow and found that the effects of climate were greater than those due to land-use change. Building on that work, Cheng et al. (2017) used SWAT to investigate the ability of stormwater detention to mitigate the effects of climate change on high flows for the Charles River watershed in Massachusetts.

In this work, we use SWAT to examine the effects of plausible, future land-use scenarios on water-related ecosystem services for two watersheds in New England under both a historical and potential future climate. The land-use scenarios were co-developed with scientists and a range of stakeholders as part of the New England Landscape Futures (NELF) project, a large research network designed to integrate diverse modes of knowledge and create a shared understanding of how the future may unfold (McBride et al., 2019). Like all scenarios, the NELF scenarios are not intended as forecasts or predictions; instead, they explore multiple hypothetical futures in a way that recognizes the irreducible uncertainty and unpredictability of complex systems (Thompson et al., 2012). Co-designing scenarios increases the range of viewpoints included in the process and is widely credited with enhancing the relevance, credibility, and salience of outcomes (Cash et al., 2003). Participatory development of land-use scenarios is particularly useful in landscapes such as New England where change is driven by the behaviors and decisions of thousands of independent land owners rather than by a central decision-making authority. Throughout this paper, we use the term “scenarios” to refer to the stakeholder-informed future landscapes, and we use the term “simulations” to refer to the combinations of climate-watershed-landscape used in our analyses.

In New England, where precipitation is abundant and consistent throughout the year, stakeholders expressed that the primary water-quantity issues of concern are related to stormwater, peak streamflow, and flooding. Consequently, this work focuses on effects of land use on storm runoff and high flows. The intent is to reveal the magnitude and robustness of potential effects due to plausible changes to the landscape. This work can provide one piece of a more holistic and comprehensive assessment of ecosystem services across these land-use scenarios (e.g., Thompson et al., 2014).

## 2 Methods

### 2.1 Land-cover scenarios for New England in 2060

McBride et al. (2017) and McBride et al. (2019) describe NELF’s participatory process for co-developing scenarios of future land cover in New England in 2060. In brief, four narrative land-use scenarios were co-designed in context with a “Recent Trends” scenario using a scenario development process that engaged over 150 stakeholders (e.g., conservationists, planners, resource managers, land owners, scientists, etc.) from throughout the region. The scenarios were created using the Intuitive Logics approach, a structured process in which participants develop plausible storylines describing a set of distinct alternative futures (Schwarz, 1991). The NELF participants used this process to construct four scenarios – Go It Alone (GA), Connected Communities (CC), Yankee Cosmopolitan (YC), and Growing Global (GG) – characterized by extreme states of two driver variables: (1) low to high natural resource planning and innovation and (2) local to global socio-economic connectedness (Table 1), which they determined to be among the most uncertain and potentially impactful for the region. Storylines for each scenario are provided in Table 1, which is adapted from the detailed narratives available in Fallon Lambert et al. (2018).

**Table 1.**
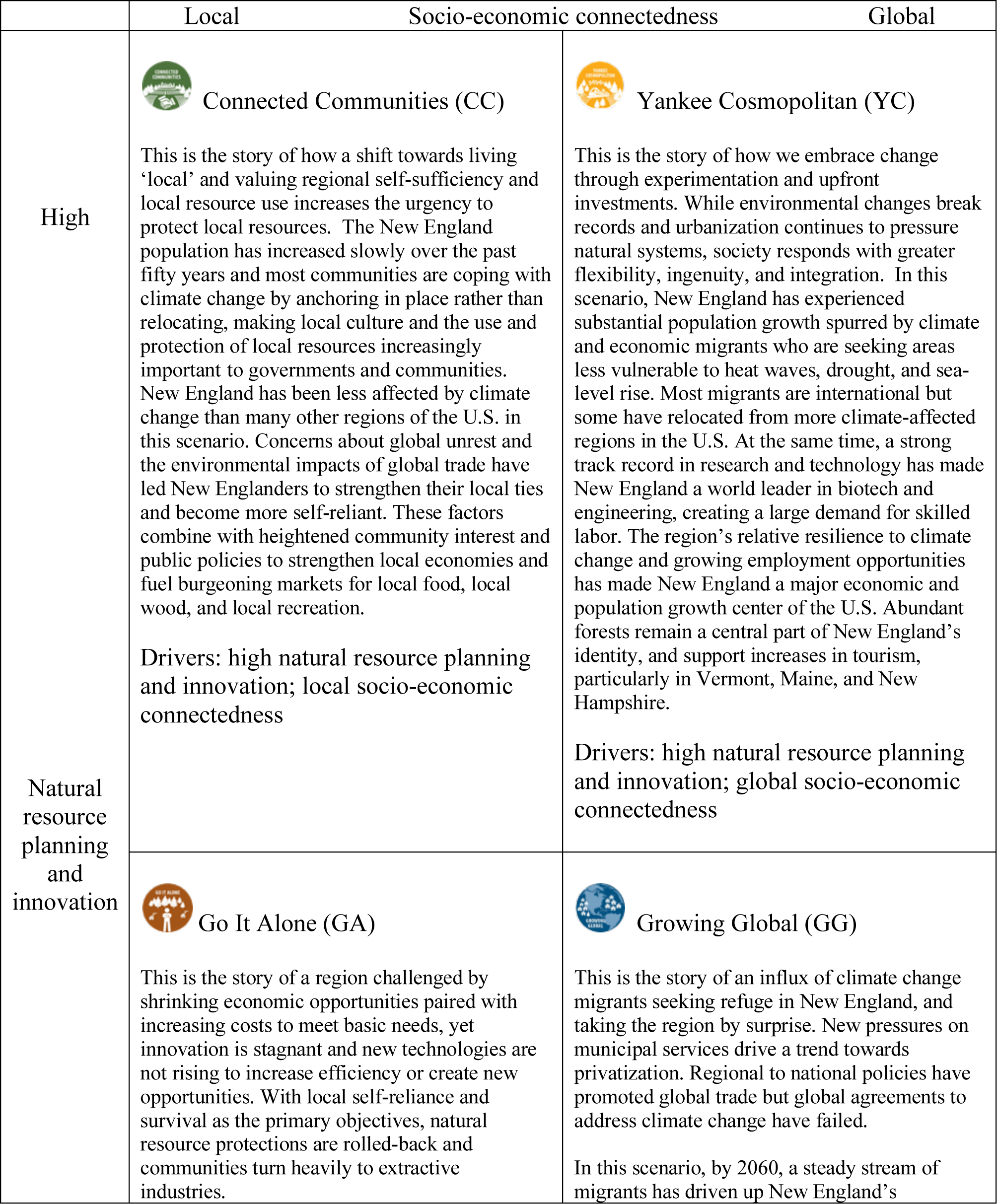
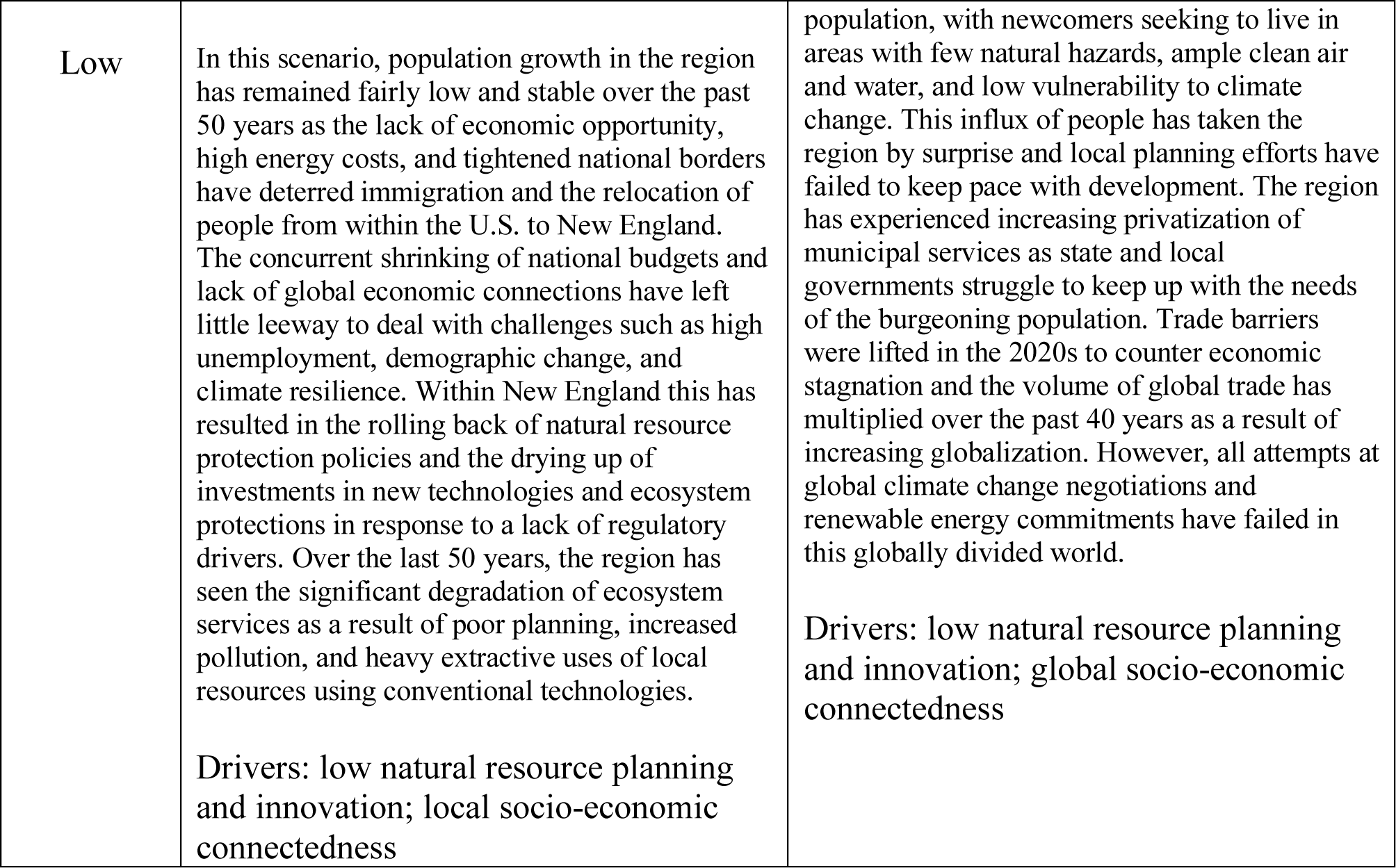
Storylines for New England Landscape Futures in 2060. Scenario icons and descriptions are adapted with permission from Fallon Lambert et al. (2018).

Land uses for the scenarios were simulated using the cellular land-cover-change model, Dinamica EGO v.2.4.1 (Soares-Filho et al. 2009; Soares-Filho et al., 2013), using a process that iterated between modelers and stakeholders to ensure that the resulting maps accurately represented the stakeholders’ intent (Thompson et al., 2017). The 50-year simulations have 30-m resolution and span the years 2010 to 2060 in ten-year time steps. Land cover varies across five classes: High Density Development, Low Density Development, Forest, Agriculture, and Legally Protected Land (e.g., conservation easements). Other land-cover classes, such as water, were held constant throughout the simulations. For the Recent Trends scenario, the rate and spatial patterns of land-cover transitions were based on observed changes in classified Landsat data between 1990 and 2010 (Olofsson et al. 2016; Thompson et al., 2017).

### 2.2 Study watersheds – Cocheco River and Charles River

To investigate the effects of these plausible landscape scenarios (Table 1) on streamflow, we selected the Cocheco River watershed, defined by USGS gage 01072800, and the Charles River watershed, defined by USGS gage 01104500 (Figure 1). The Cocheco River watershed in southeastern New Hampshire was selected because it is in one of the most rapidly urbanizing parts of New England. The watershed has an area of 207 km^2^, and the main channel is 34 km in length and drops 170 m in elevation from the headwaters to the gage at an elevation of 36.2 m. Average annual precipitation is 1059 mm/year, and average streamflow is 3.14 cms, equivalent to 479 mm/year. The Charles River flows through some of the most densely populated parts of New England, and a SWAT model had previously been calibrated to study this watershed (Cheng et al., 2017). The watershed has an area of 648 km^2^, and it is flatter and more developed than the Cocheco watershed (Figure 1). The outlet elevation is 6.10 m, and the main channel drops only 101 m over its 108-km length. Average annual precipitation is 1111 mm/year, and average streamflow is 8.01 cms, equivalent to 389 mm/year. Figures 2 and 3 display the land uses across the Cocheco River and Charles River watersheds for the landscape scenarios described above. Table 2 reports the fraction of each land-use type within the watersheds.

**Figure 1.**
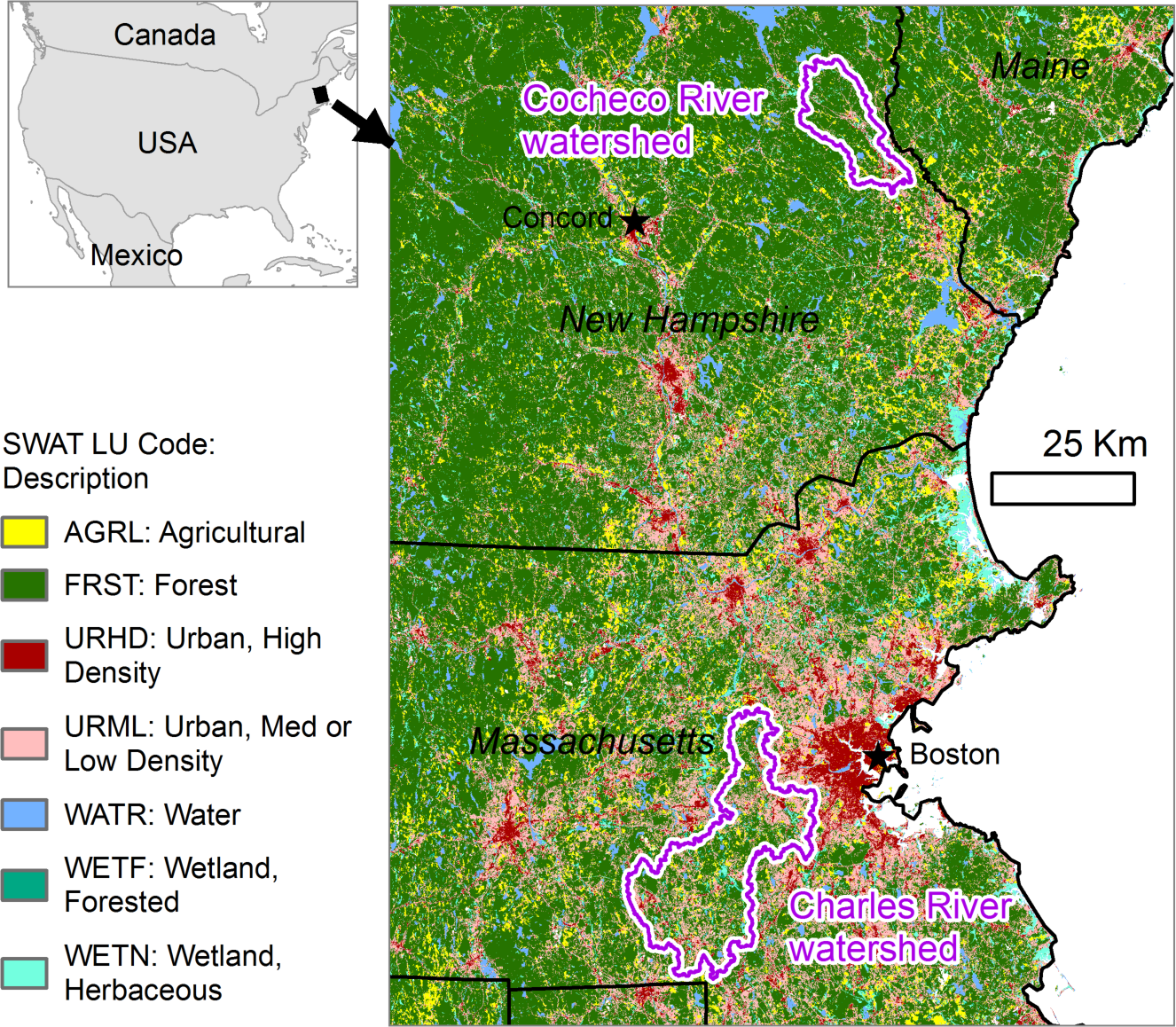
Locations of the Cocheco River watershed and the Charles River watershed with land covers from 2010. [color]

**Figure 2.**
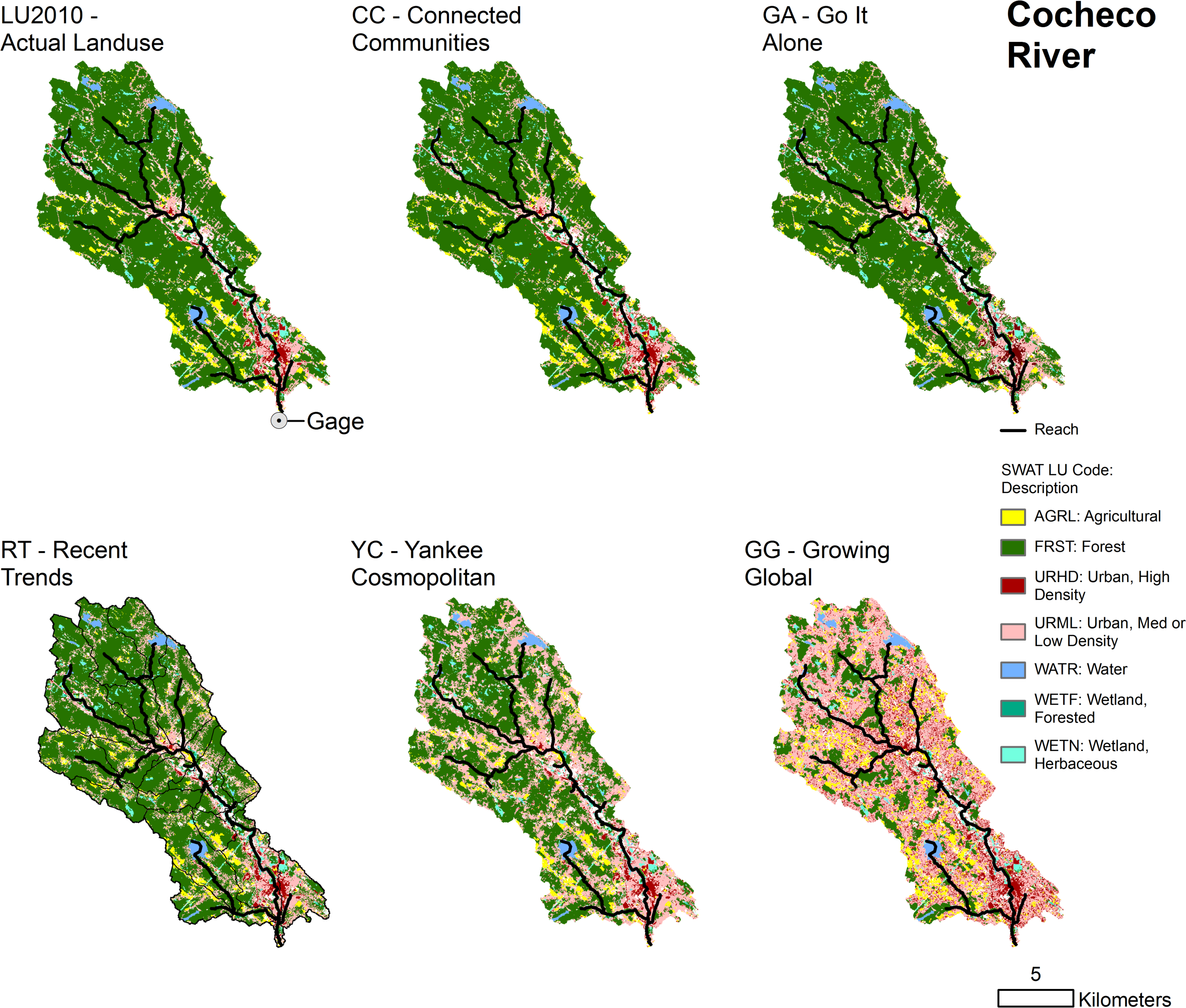
Land use for the Cocheco River watershed. LU2010 indicates land use in 2010 from the NLCD. Recent Trends, Connected Communities, Yankee Cosmopolitan, Go It Alone, and Growing Global represent plausible land-use scenarios in 2060. [color]

**Figure 3.**
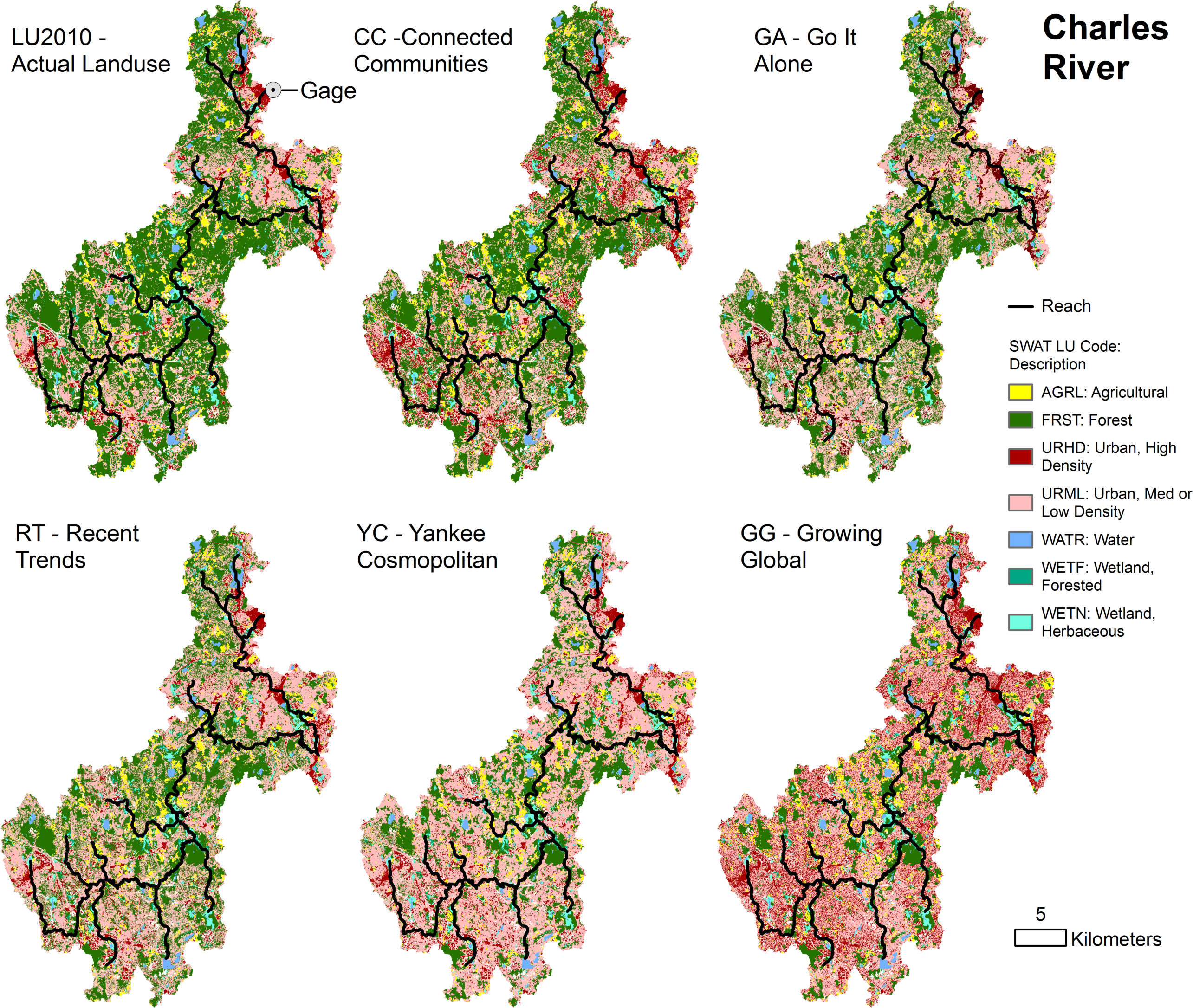
Land use for the Charles River watershed. LU2010 indicates land use in 2010 from the NLCD. Recent Trends, Connected Communities, Yankee Cosmopolitan, Go It Alone, and Growing Global represent plausible land-use scenarios in 2060. [color]

**Table 2.** Fractional land use across scenarios and watersheds.

| Watershed | Land-use scenario | % Urban | % Forest | % Agriculture |
| --- | --- | --- | --- | --- |
| Cocheco | 2010 | 14.3 | 75.5 | 4.4 |
|  | RT | 22.8 | 66.7 | 4.7 |
|  | GA | 16.1 | 73.8 | 4.3 |
|  | CC | 14.6 | 74.3 | 5.3 |
|  | YC | 39.6 | 49.9 | 4.7 |
|  | GG | 65.7 | 18.6 | 9.8 |
| Charles | 2010 | 35.2 | 50.6 | 6.2 |
|  | RT | 50.7 | 35.2 | 6.1 |
|  | GA | 47.7 | 38.4 | 5.8 |
|  | CC | 41.8 | 43.3 | 6.9 |
|  | YC | 63.8 | 22.8 | 5.4 |
|  | GG | 68.9 | 16.2 | 6.8 |

### 2.3 Hydrologic model – SWAT

This project employed the Soil and Water Assessment Tool (SWAT; Arnold et al., 1998; Nietsch et al., 2011) to represent the effects of land-cover differences on hydrology. SWAT is a well-known and proven process-based model that represents weather, hydrology, growth and seasonality of vegetation, and landscape management practices. It operates with a daily time step, and space is represented in a semi-distributed way. Within a watershed, sub-basins are linked via a stream network, and each sub-basin is represented by a collection of Hydrologic Response Units (HRUs). Each HRU comprises a particular combination of soil, slope, land use, and land management. Within a sub-basin, HRUs are not represented explicitly in space and do not interact with each other, and water-balance equations are solved within each HRU. Incoming precipitation is partitioned among canopy interception, storm runoff, and storage in the soil. Soil water then contributes to lateral subsurface flow, groundwater return flow, and deep recharge.

In SWAT, ecosystem services related to hydrology and streamflow are affected by a few parameters related to land use, including

- The curve number, the parameter in the Curve Number Method for estimating storm runoff (USDA, 2004), which depends on soil and land use;
- The fraction of impervious area and the fraction of impervious area that is directly connected to storm sewer infrastructure; these parameters affect the aggregated curve number for urban areas;
- Vegetation, which affects the seasonality and magnitude of evapotranspiration.

Across different landscape scenarios, the weather, topography, soils, and soil-related parameters are held constant.

#### 2.3.1 Weather forcing

Simulations were run in SWAT for twenty-year periods for both historical weather and a future climate. The first three years of all simulations were used as a warm-up period and were not used in subsequent analyses. The years were selected so that the final simulated years coincided with the years of the landcover datasets plus the eight years before and the eight years after (2002-2017 for the historical weather and 2052-2067 for the simulated future climate).

Data for the historical weather came from the National Oceanic and Atmospheric Administration’s Climate Data Online Search webtool (NOAA, 2018). We used data from the Rochester Skyhaven Airport (054791) weather station for the Cocheco River watershed and the Boston (14739) weather station for the Charles River watershed. Precipitation and temperature data for a possible future climate were obtained from the USGS Geo Data Portal Bias Corrected Constructed Analogs V2 Daily Climate Projections dataset (USGS, 2018). The spatially and temporally downscaled LOCA CMIP5 CCSM4 RCP 8.5 dataset has among the highest temperature correlations with observed data (Kumar et al., 2013) and performs well with comparisons to historical and paleo climate data (Sillmann et al., 2013).

Precipitation and temperature data were used with the weather generator in SWAT (using the WGEN_US_FirstOrder database) to simulate additional weather parameters, including relative humidity, wind speed, and solar radiation. The Penman-Monteith method was used to estimate potential evapotranspiration (Arnold et al., 2012).

#### 2.3.2 Landscape features and watershed discretization

Land-cover maps were derived from the NELF scenarios (McBride et al., 2017; McBride et al., 2019; Plisinski et al., 2017). While the NELF scenarios combined water with wetlands and considered swamps to be forests, we separated water and wetlands, and accounted for herbaceous wetlands and swamps explicitly by extracting those land-cover types from the National Land Cover Database (NLCD; Homer et al., 2015) and imposing them on the NELF scenarios. Soil data were obtained from the SSURGO database (USDA, 2014). Three slope classes were calculated for each watershed using natural class breaks; breakpoints of 5.7% and 14.1% were used for the Cocheco River watershed and 4.6% and 11.5% for the Charles River watershed. To better represent the small land-use patches that are typical of the New England landscape, we did not merge smaller HRUs with larger neighbors, as is sometimes done.

Land-cover differences in SWAT manifest predominantly as differences in plant growth and evapotranspiration and in the generation of storm runoff via the curve number (Arnold et al., 2012). We chose curve numbers (CN2) to reflect conditions in New England. Because there is very little woodland pasturing in New England, we changed the CN2 values for generic forest (FRST) from the default values in SWAT, which would be appropriate in forests subject to grazing by livestock (“fair” condition), to those for forests without livestock grazing (“good” condition). CN2 values for forest were 5, 55, 70, 77 for soil hydric classes A through D, respectively.

The New England Landscape Futures use a single designation for all agricultural land, and we do the same by using the generic agriculture land-cover (AGRL). The default curve numbers for AGRL in SWAT are appropriate for farmland dominated by corn or row crops, while New England farms are primarily pasture and hay fields. Therefore, we updated this parameter by using county-level data from the United States Agricultural Census (USDA, 2018) to determine an area-weighted curve number based on the actual agricultural types. Resulting curve numbers for our agricultural land use are 42.3, 65.1, 76.2, and 82.1 for soil hydric classes A through D, respectively. Other parameters in SWAT’s vegetation database (plant.dat) were not changed.

Urban areas in the NELF scenarios are designated as either “high-density development” or “low-density development.” We consider these two classes to be analogous to Urban Residential High Density (URHD) and Urban Residential Medium/Low Density (URML), respectively, in SWAT. We calculated the fraction of impervious surface for all of New England by overlaying the NLCD urban landcover types (Homer et al., 2015) on the NLCD 2011 Percent Developed Imperviousness GIS layer (Xian et al., 2011) and calculating separate area-weighted averages for URHD (consisting of the NLCD “Developed, High Intensity”) and URML (consisting of NLCD “Developed, Open Space”, “Developed, Low Intensity”, and “Developed Medium Intensity”). This resulted in 88.9% impervious for URHD and 27.5% impervious for URML in our simulations. For all scenarios except Connected Communities and Yankee Cosmopolitan, the fractions of connected impervious area (i.e., the impervious area that is directly connected to storm sewers) were left at SWAT’s default values of 44% and 17%, respectively, for URHD and URML. For Connected Communities and Yankee Cosmopolitan, those numbers were halved to 22% and 8.5% to represent natural-resource innovation (Table 1) and the implementation of green infrastructure, such as bioswales and rain gardens. CN2 values for the pervious portions of these urban areas were set to 39, 61, 74, 80 for hydric classes A through D, respectively, since the pervious portions of New England’s urban areas are usually grass-covered lawns with greater than 75% grass cover (Arnold et al., 2012). No other values in the urban.dat file were changed.

#### 2.3.3 Calibration and model performance under historic conditions

To increase model performance and accuracy, parameters that were unrelated to land cover within the SWAT model were calibrated by matching simulated streamflow to observed streamflow under current land use. Model parameters were calibrated separately for each watershed using observed flow for the years 2002-2011 and validated using the observed flow from 2012-2017. Parameters that were explicitly related to land use, such as the curve number and vegetation-related parameters, were not included in calibration, since they were our driver variables of interest. We used a semi-automated approach with the SWAT Calibration and Uncertainty Program (SWAT-CUP) using the SUFI-2 optimization method (Abbaspour, 2015). Starting values for our calibration were either the default values in SWAT or the calibrated results from an earlier study on the Charles River (Cheng, et al., 2017). We used the Nash-Sutcliffe efficiency, percent bias, and the ratio of the root-mean-square error to the standard deviation of the streamflow observations (RSR) as metrics of goodness-of-fit. Calibration continued until none of the metrics improved by more than 5% over the previous iteration.

The final model for the Cocheco River had a NSE of 0.58, RSR of 0.64, and percent bias of −13.6% for the calibration period. The model for the Charles River had values of 0.74, 0.51, and 1.2%, respectively. Moriasi et al. (2007) suggest that a model can be viewed as satisfactory if the NSE value is greater than 0.50, the RSR is less than 0.70, and the percent bias is less than plus or minus 25%; the calibrated models for both rivers were deemed satisfactory. For the validation period, the Cocheco River had a NSE of 0.49, RSR of 0.72, and percent bias of −19.5%, and the Charles River had values of 0.74, 0.51, and 23.3% respectively. Final parameters and goodness-of-fit metrics are shown in Table 3.

**Table 3.** Calibrated parameters for SWAT models of the Cocheco River and Charles River watersheds.

| Description | SWAT file | Cocheco | Charles |
| --- | --- | --- | --- |
| Soil evaporation compensation factor | .bsn | 0.84 | 0.99 |
| Surface runoff lag coefficient | .bsn | 3.94 | 4.62 |
| Fraction of transmission loss from main channel that enter deep aquifer | .bsn | 0.01 | 0.01 |
| Plant uptake compensation factor | .bsn | 0.69 | 0.95 |
| Baseflow alpha factor (1/days) | .gw | 0.19 | 0.22 |
| Groundwater delay time (days) | .gw | 32.16 | 36.37 |
| Threshold depth in the shallow aquifer required for return flow (mm H<sub>2</sub>O) | .gw | 972.19 | 1150.78 |
| Groundwater "revap" coefficient | .gw | 0.06 | 0.08 |
| Deep aquifer percolation fraction | .gw | 0.09 | 0.08 |
| Threshold depth in the shallow aquifer for "revap" or percolation to the deep aquifer (mm H<sub>2</sub>O) | .gw | 1027.11 | 534.91 |
| Soil evaporation compensation factor for HRUs | .hru | 0.98 | 1.00 |
| Plant uptake compensation factor | .hru | 0.78 | 0.09 |
| Baseflow alpha factor for bank storage (days) | .rte | 0.81 | 0.70 |
| Effective hydraulic conductivity in main channel alluvium (mm/hr) | .rte | 242.34 | 405.11 |
| Available water content of the soil (mm H<sub>2</sub>O/mm soil) for hydric class A | .sol | 0.22 | 0.20 |
| Available water content of the soil (mm H<sub>2</sub>O/mm soil) for hydric class B | .sol | 0.30 | 0.30 |
| Available water content of the soil (mm H<sub>2</sub>O/mm soil) for hydric class C | .sol | 0.19 | 0.08 |
| Available water content of the soil (mm H<sub>2</sub>O/mm soil) for hydric class D | .sol | 0.15 | 0.16 |
| Saturated hydraulic conductivity (mm/hr) for hydric class A | .sol | 236.58 | 65.47 |
| Saturated hydraulic conductivity (mm/hr) for hydric class B | .sol | 391.94 | 492.62 |
| Saturated hydraulic conductivity (mm/hr) for hydric class C | .sol | 259.36 | 252.56 |
| Saturated hydraulic conductivity (mm/hr) for hydric class D | .sol | 321.95 | 343.35 |
| Effective hydraulic conductivity in tributary channel alluvium (mm/hr) | .sub | 170.37 | 330.13 |

### 2.4 Streamflow metrics of interest

Across scenarios and climates, we consider two metrics of hydrologic regulation. The first is the water balance – the partitioning of precipitation among evapotranspiration, storm runoff, and baseflow. Runoff and baseflow together constitute streamflow; storm runoff is the rapid response to precipitation events, whereas baseflow represents the slower component of streamflow driven by seasonal and interannual variability. The second metric is the annual maximum daily flow. While true peak flows may be short-lived phenomena – on the scale of minutes to hours – the annual maximum daily flow nonetheless provides an indication of the potential for flooding and associated damage.

## 3 Results

### 3.1 Water balance

Across the simulations, land use has little effect on the average partitioning of precipitation between evapotranspiration and streamflow (Figure 4). Under historic weather, simulated evapotranspiration is 44-45% of precipitation in the Cocheco River watershed and 46-48% of precipitation in the Charles River watershed with little variation among land-use scenarios (Table 5). For the future climate, annual precipitation increases from 1059 mm to 1194 mm in the Cocheco River watershed and 1111 mm to 1345 mm in the Charles River watershed, and potential evaporation decreases (Table 5). As a result, evaporation represents a smaller fraction (35%-36%) of precipitation for the simulations with a future climate.

**Figure 4.**
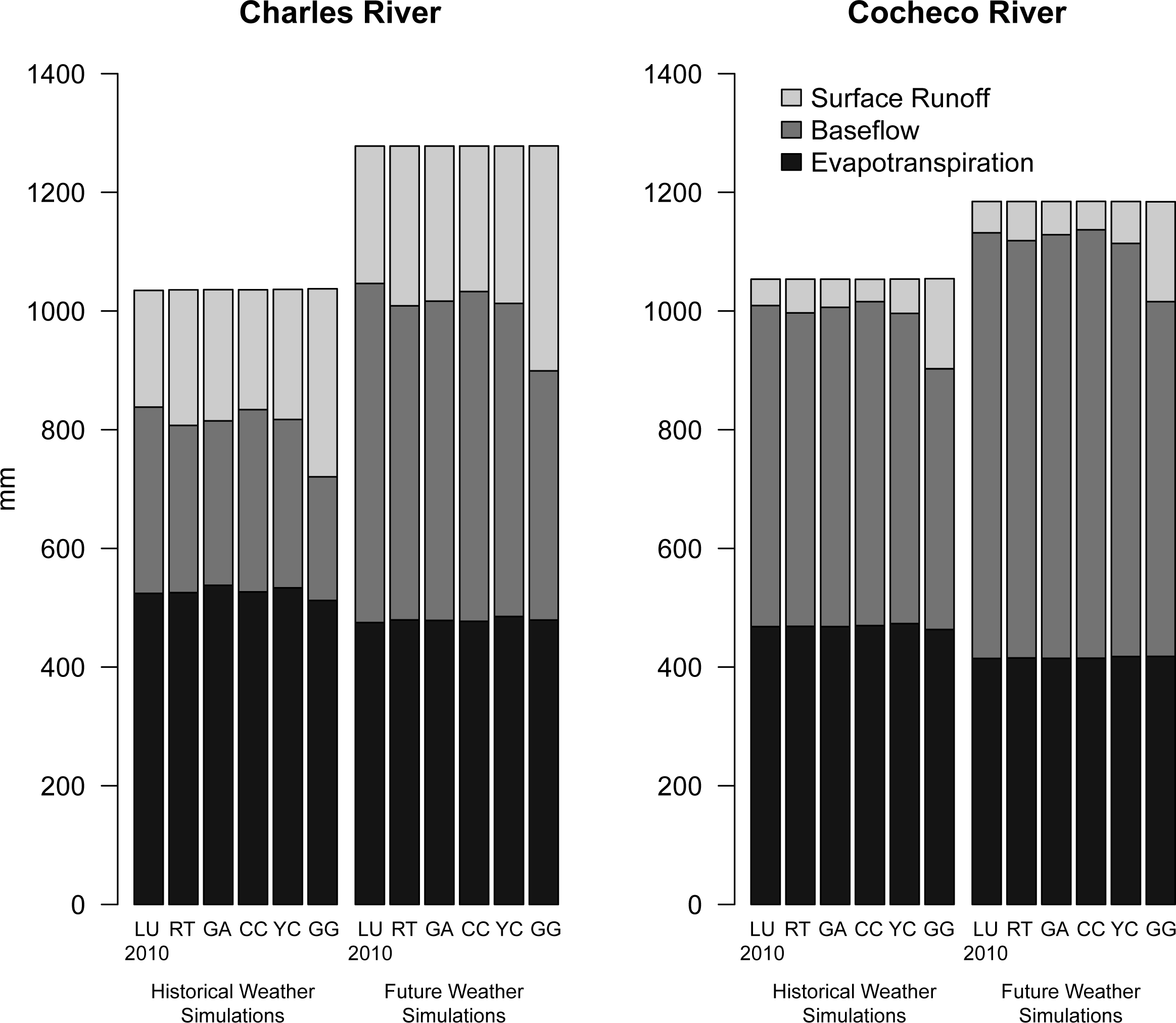
Average water balances for the Charles River and Cocheco River watersheds across scenarios and climates.

**Table 4.** Goodness-of-fit for SWAT models of the Charles River and Cocheco River watersheds. NSE is the Nash-Sutcliffe efficiency, and RSR is the ratio of the root-mean-squared error to the standard deviation of the observations. All goodness-of-fit statistics were calculated in R with the hydroGOF package (Zambrano-Bigiarini, 2017).

| Watershed | Description | Years | NSE | Bias | RSR |
| --- | --- | --- | --- | --- | --- |
| Cocheco | calibration | 2002–2011 | 0.58 | –13.6% | 0.64 |
|  | validation | 2012–2017 | 0.49 | –19.5% | 0.72 |
|  | overall | 2002–2017 | 0.58 | –15.3% | 0.65 |
| Charles | calibration | 2002–2011 | 0.74 | 1.2% | 0.51 |
|  | validation | 2012–2017 | 0.74 | 23.3% | 0.51 |
|  | overall | 2002–2017 | 0.75 | 7.2% | 0.5 |

**Table 5.**
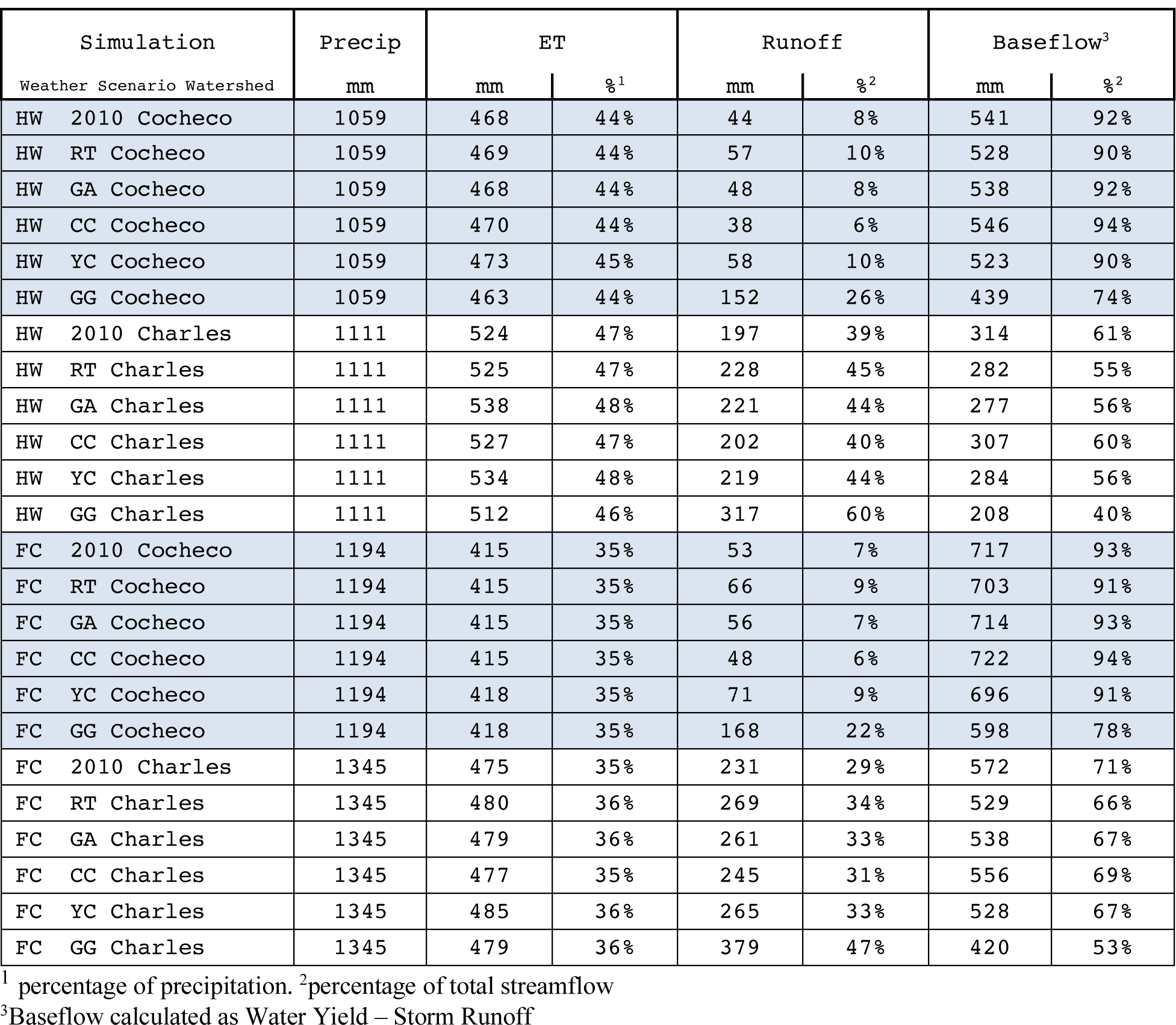
Average annual fluxes across simulations. HW and FC indicate simulations under historic weather and a future climate, respectively. Scenarios are denoted as follows: 2010 – historic land use in 2010, RT – Recent Trends, GA – Go It Alone, CC – Connected Communities, YC – Yankee Cosmopolitan, GG – Growing Global.

| Simulation |  |  | Precip |  | ET |  | Runoff |  | Baseflow<sup>3</sup> |  |
| --- | --- | --- | --- | --- | --- | --- | --- | --- | --- | --- |
| Weather | Scenario | Watershed | mm |  | mm | %<sup>1</sup> | mm | %<sup>2</sup> | mm | %<sup>2</sup> |
| HW | 2010 | Cocheco | 1059 |  | 468 | 44% | 44 | 8% | 541 | 92% |
| HW | RT | Cocheco | 1059 |  | 469 | 44% | 57 | 10% | 528 | 90% |
| HW | GA | Cocheco | 1059 |  | 468 | 44% | 48 | 8% | 538 | 92% |
| HW | CC | Cocheco | 1059 |  | 470 | 44% | 38 | 6% | 546 | 94% |
| HW | YC | Cocheco | 1059 |  | 473 | 45% | 58 | 10% | 523 | 90% |
| HW | GG | Cocheco | 1059 |  | 463 | 44% | 152 | 26% | 439 | 74% |
| HW | 2010 | Charles | 1111 |  | 524 | 47% | 197 | 39% | 314 | 61% |
| HW | RT | Charles | 1111 |  | 525 | 47% | 228 | 45% | 282 | 55% |
| HW | GA | Charles | 1111 |  | 538 | 48% | 221 | 44% | 277 | 56% |
| HW | CC | Charles | 1111 |  | 527 | 47% | 202 | 40% | 307 | 60% |
| HW | YC | Charles | 1111 |  | 534 | 48% | 219 | 44% | 284 | 56% |
| HW | GG | Charles | 1111 |  | 512 | 46% | 317 | 60% | 208 | 40% |
| FC | 2010 | Cocheco | 1194 |  | 415 | 35% | 53 | 7% | 717 | 93% |
| FC | RT | Cocheco | 1194 |  | 415 | 35% | 66 | 9% | 703 | 91% |
| FC | GA | Cocheco | 1194 |  | 415 | 35% | 56 | 7% | 714 | 93% |
| FC | CC | Cocheco | 1194 |  | 415 | 35% | 48 | 6% | 722 | 94% |
| FC | YC | Cocheco | 1194 |  | 418 | 35% | 71 | 9% | 696 | 91% |
| FC | GG | Cocheco | 1194 |  | 418 | 35% | 168 | 22% | 598 | 78% |
| FC | 2010 | Charles | 1345 |  | 475 | 35% | 231 | 29% | 572 | 71% |
| FC | RT | Charles | 1345 |  | 480 | 36% | 269 | 34% | 529 | 66% |
| FC | GA | Charles | 1345 |  | 479 | 36% | 261 | 33% | 538 | 67% |
| FC | CC | Charles | 1345 |  | 477 | 35% | 245 | 31% | 556 | 69% |
| FC | YC | Charles | 1345 |  | 485 | 36% | 265 | 33% | 528 | 67% |
| FC | GG | Charles | 1345 |  | 479 | 36% | 379 | 47% | 420 | 53% |

<sup>1</sup> percentage of precipitation. <sup>2</sup>percentage of total streamflow
<sup>3</sup>Baseflow calculated as Water Yield – Storm Runoff

While total streamflow is nearly unchanged across the land-use scenarios, the partitioning of streamflow between baseflow and storm runoff does vary. In the Cocheco River watershed, baseflow is 90-94% of streamflow for all land-use scenarios, except Growing Global, for both historic and future weather. For Growing Global, baseflow is 74% and 78% of streamflow for historic weather and a future climate, respectively. In the more developed Charles River watershed, baseflow represents between 40% and 61% of streamflow under historic weather, with the lowest fraction associated with the Growing Global scenario (Table 5). For the future climate, both storm runoff and baseflow increase. As a fraction of streamflow, the baseflow contribution increases by approximately 10% and shows variability across scenarios similar to that under historic weather.

Seasonal water balances exhibit behavior similar to the annual water balances. Differences in land use have little effect on the partitioning of water between streamflow and evapotranspiration; rather, the effect is in the separation of streamflow into baseflow and storm runoff (Table 6 and 7). The increases in streamflow associated with a future climate vary seasonally, with large increases in autumn and winter, moderate increases in spring, and little effect in summer (Figure 5). For historic weather, streamflow during the fall and winter represents 40% and 44% of annual streamflow for the Cocheco River and Charles River watersheds, respectively. Those fractions increase to 51% and 55% under a future climate (Figure 5).

**Table 6.** Seasonal evapotranspiration (ET), storm runoff (SR), and baseflow (BF), in mm, for the Cocheco River watershed.

|  | Historic Weather |  |  |  |  |  | Future Climate |  |  |  |  |  |
| --- | --- | --- | --- | --- | --- | --- | --- | --- | --- | --- | --- | --- |
|  | 2010 Land Use |  |  | Growing Global |  |  | 2010 Land Use |  |  | Growing Global |  |  |
|  | ET | SR | BF | ET | SR | BF | ET | SR | BF | ET | SR | BF |
| Fall | 93 | 11 | 126 | 92 | 41 | 105 | 84 | 11 | 155 | 82 | 36 | 125 |
| Winter | 29 | 6 | 94 | 27 | 22 | 73 | 20 | 19 | 203 | 18 | 52 | 161 |
| Spring | 129 | 19 | 196 | 122 | 56 | 155 | 118 | 18 | 247 | 112 | 60 | 209 |
| Summer | 218 | 8 | 125 | 222 | 34 | 107 | 193 | 5 | 112 | 206 | 21 | 103 |

**Table 7.** Seasonal evapotranspiration (ET), storm runoff (SR), and baseflow (BF), in mm, for the Charles River watershed.

|  | Historic Weather |  |  |  |  |  | Future Climate |  |  |  |  |  |
| --- | --- | --- | --- | --- | --- | --- | --- | --- | --- | --- | --- | --- |
|  | 2010 Land Use |  |  | Growing Global |  |  | 2010 Land Use |  |  | Growing Global |  |  |
|  | ET | SR | BF | ET | SR | BF | ET | SR | BF | ET | SR | BF |
| Fall | 113 | 42 | 42 | 115 | 70 | 25 | 94 | 59 | 115 | 96 | 95 | 76 |
| Winter | 43 | 43 | 100 | 42 | 69 | 70 | 34 | 72 | 192 | 34 | 115 | 144 |
| Spring | 129 | 66 | 121 | 124 | 102 | 84 | 130 | 60 | 186 | 128 | 99 | 145 |
| Summer | 239 | 45 | 51 | 232 | 76 | 30 | 217 | 41 | 79 | 221 | 71 | 55 |

**Figure 5.**
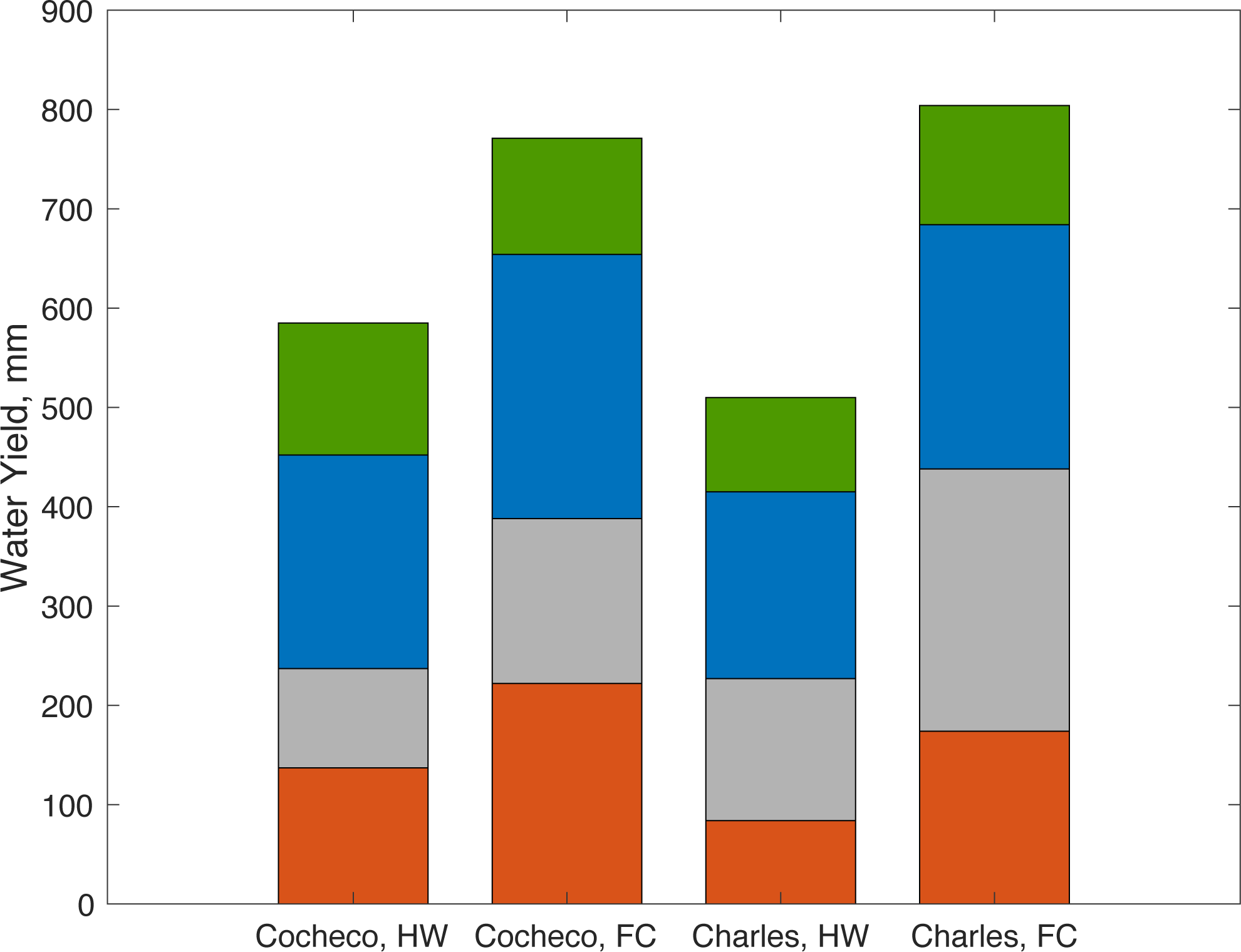
Seasonal water yield for the Cocheco and Charles River watersheds under for land use from 2010 and historic weather (HW) and a future climate (FC). Seasons are represented by colors, from the bottom: orange – autumn (SON); gray – winter (DJF); blue – spring (MAM); green – summer (JJA). [color]

### 3.2 Changes in magnitude of annual maximum daily flow

The annual maximum daily flow (AMDF) exhibits significant year-to-year variability due to variability in weather and precipitation. Under historic weather and land use, simulated AMDFs range from 10.7 to 78.2 m^3^/s (equivalent to 4.5 to 32.7 mm/day) for the Cocheco River watershed and 15.5 to 94.0 m^3^/s (2.1 to 12.5 mm/day) for the Charles River watershed. Due to this year-to-year variability, a paired comparison was used to quantify the effect of land use on AMDF. For each year of simulation, the difference in AMDF between each future land-use scenario and the land use from 2010 indicates the effect of land-use change on high flows. Figures 6 and 7 present the average differences, along with their 95%-confidence intervals determined via 10,000 bootstrap samples, expressed as a percent of the average AMDF under land use in 2010.

**Figure 6.**
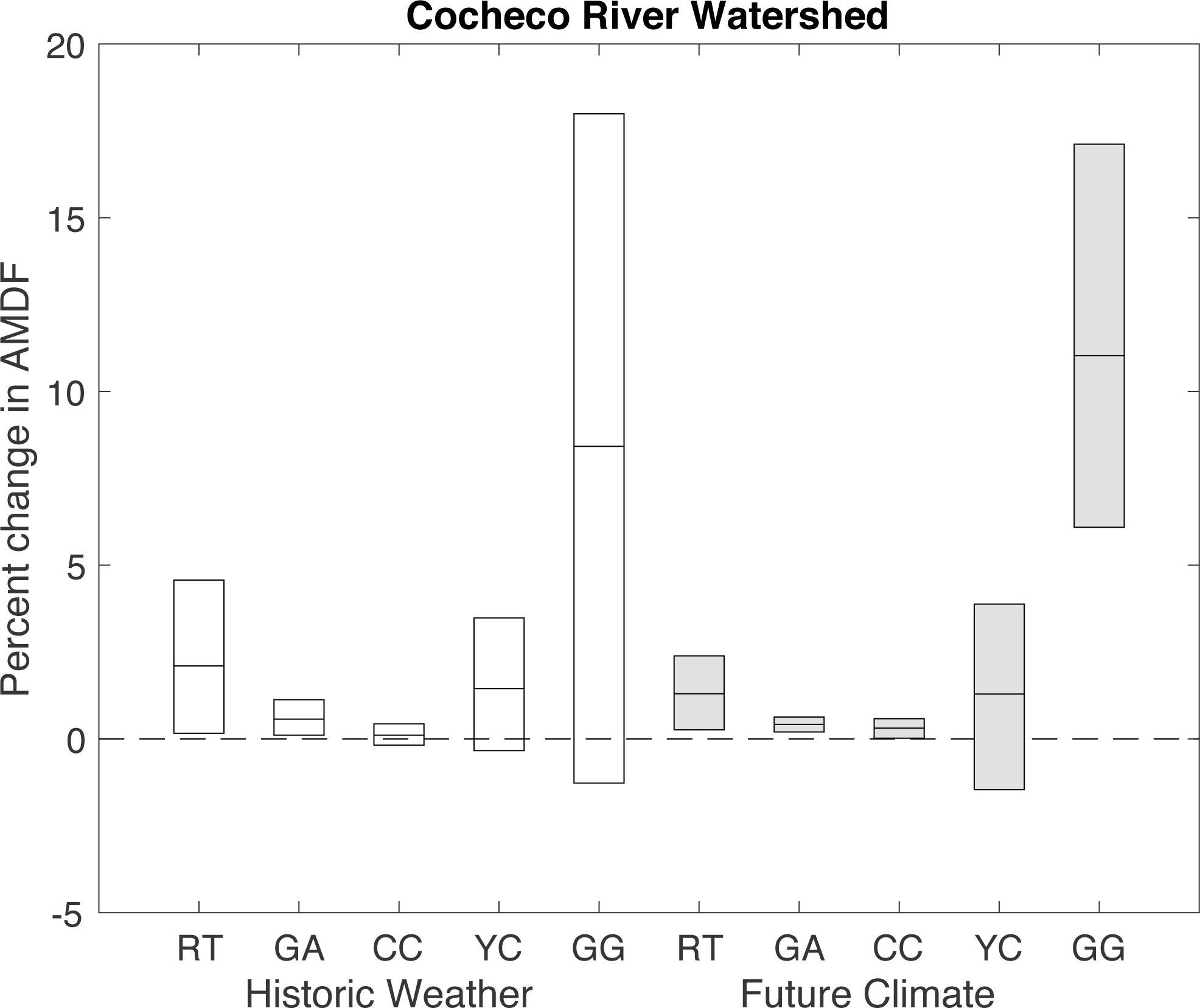
Difference in annual maximum daily flow (AMDF) between land-use scenarios and historic land use in 2010 for the Charles River watershed. Bars represent 95%-confidence limits determined via bootstrap, and the line in the middle represents the mean difference in AMDF between that scenario and land use in 2010. Scenarios are denoted as follows: RT – Recent Trends, GA – Go It Alone, CC – Connected Communities, YC – Yankee Cosmopolitan, GG – Growing Global.

**Figure 7.**
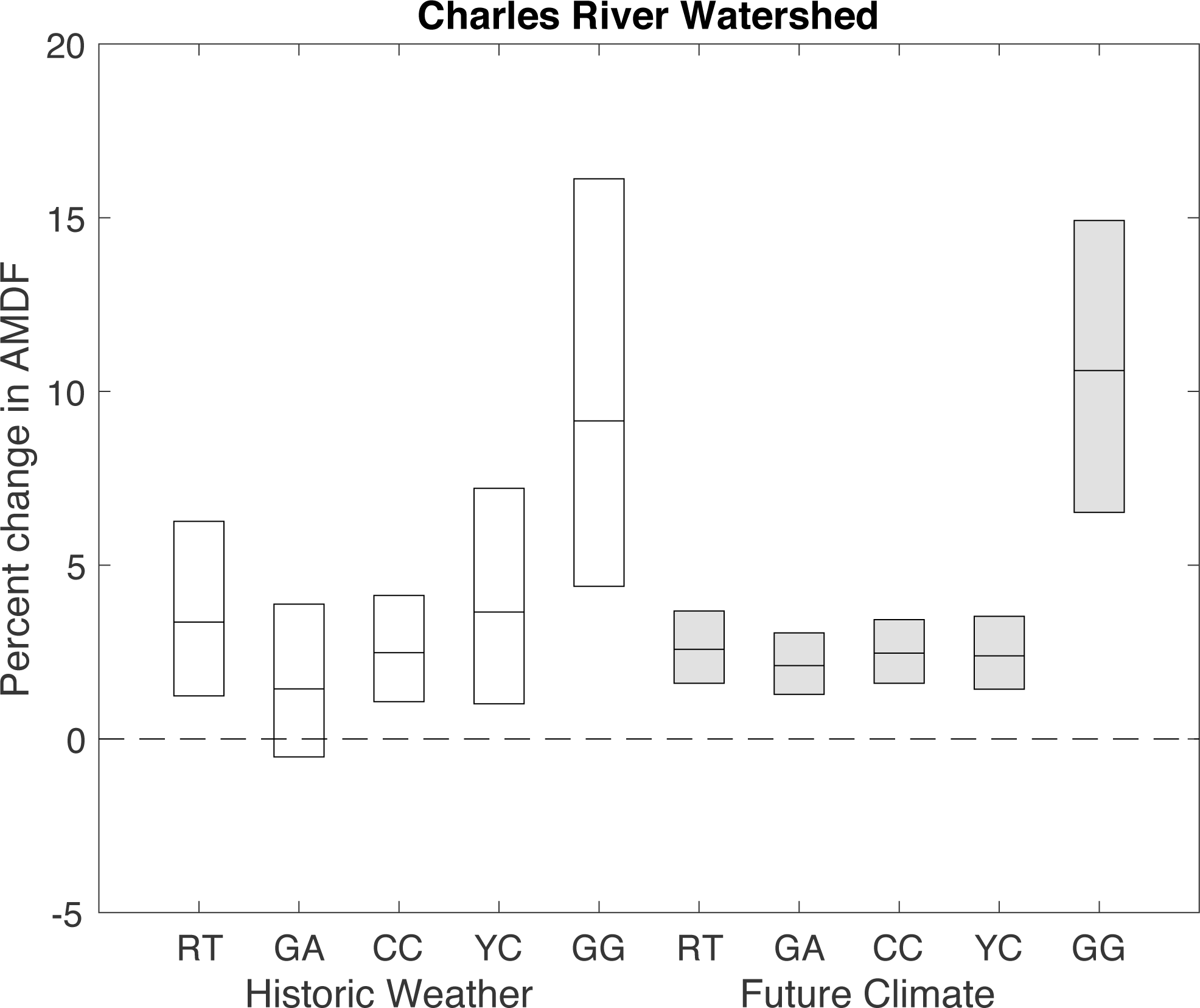
Difference in annual maximum daily flow (AMDF) between land-use scenarios and historic land use in 2010 for the Cocheco River watershed. Bars represent 95%-confidence limits determined via bootstrap, and the line in the middle represents the mean difference in AMDF between that scenario and land use in 2010. Scenarios are denoted as follows: RT – Recent Trends, GA – Go It Alone, CC – Connected Communities, YC – Yankee Cosmopolitan, GG – Growing Global.

Analysis of the difference in these flows between land use in 2010 and future scenarios indicates that land-use change could have a moderate effect on the annual maximum daily flow (Figures 6 and 7 and Table 8). Under the Growing Global scenario, the annual maximum daily flows are approximately 10% larger than those under the historic land-use scenario. This result is robust across both the Cocheco River and Charles River watersheds and both historic and future climates. Effects under other land-use scenarios are more modest, with mean values ranging from 0-4%.

**Table 8.** High-density urban area (URHD), medium/low density urban area (URML), impervious area, and average annual maximum daily flow across the simulations. Scenarios are denoted as follows: 2010 – historic land use in 2010, RT – Recent Trends, GA – Go It Alone, CC – Connected Communities, YC – Yankee Cosmopolitan, GG – Growing Global.

| Watershed | Land Use Scenario | % URHD | % URML | % Impervious | Historic Weather<br>Average Max<br>Daily Flow,<br>cms | Future Climate<br>Average Max<br>Daily Flow,<br>cms |
| --- | --- | --- | --- | --- | --- | --- |
| Cocheco | 2010 | 2.2 | 12.1 | 5.3 | 32.0 | 33.6 |
|  | RT | 2.5 | 20.3 | 7.8 | 32.7 | 34.0 |
|  | GA | 2.4 | 13.7 | 5.9 | 32.2 | 33.7 |
|  | CC | 2.9 | 11.7 | 5.8 | 32.1 | 33.7 |
|  | YC | 3.4 | 36.2 | 13.0 | 32.5 | 34.0 |
|  | GG | 14.6 | 51.1 | 27.0 | 34.7 | 37.3 |
| Charles | 2010 | 5.5 | 29.7 | 13.1 | 40.4 | 47.3 |
|  | RT | 7.5 | 43.2 | 18.5 | 41.8 | 48.5 |
|  | GA | 7.0 | 40.8 | 17.4 | 41.0 | 48.3 |
|  | CC | 11.0 | 30.8 | 18.3 | 41.4 | 48.5 |
|  | YC | 10.0 | 53.9 | 23.7 | 41.9 | 48.4 |
|  | GG | 24.4 | 44.5 | 34.0 | 44.2 | 52.3 |

While the AMDFs increase with increasing urbanization, the relationship depends on the nature of the urbanization – whether high density or medium or low density – and the associated increases in the fraction of impervious area (Figure 8). For example, while total urban area is greater for both the Recent Trends and Go-It-Alone scenarios than for Connected Communities (Table 2), the Connected Communities scenario has a higher proportion of high-density development, and a comparable fraction of total impervious area (Table 8 and Figure 8). The incorporation of green infrastructure, manifest as a lower fraction of directly connected impervious area in the Yankee Cosmopolitan and Connected Communities scenarios, mitigates the effect of urbanization on AMDF only slightly.

**Figure 8.**
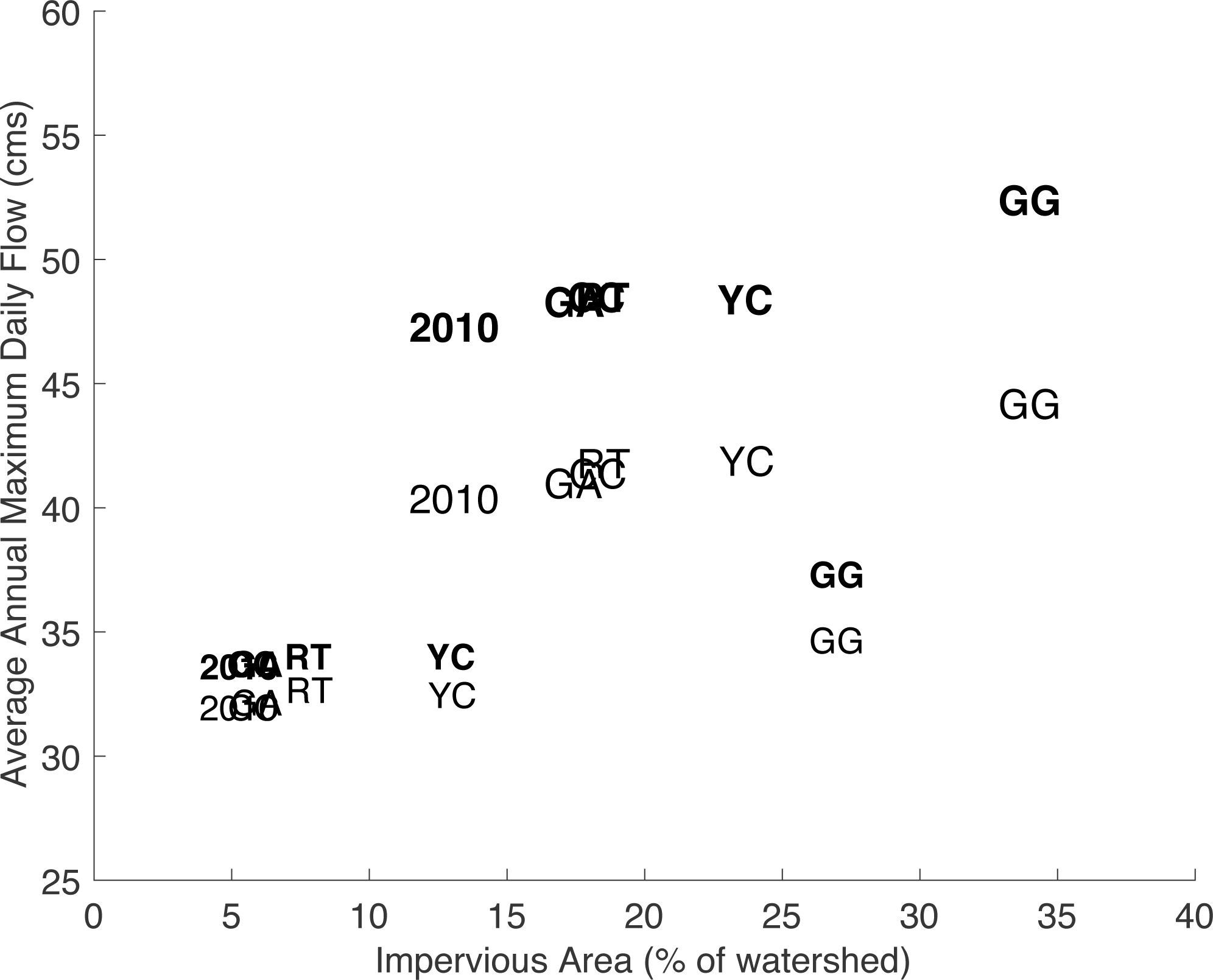
Average annual maximum daily flow increases with impervious area. Land-use scenarios are indicated by letter codes: 2010 – historic land use in 2010, RT – Recent Trends, GA – Go It Alone, CC – Connected Communities, YC – Yankee Cosmopolitan, GG – Growing Global. Bold indicates simulations for a future climate and normal font represents historic weather. The larger font (and higher magnitude flows) are for the Charles River watershed, and the smaller font is for the Cocheco River watershed. For the Charles River watershed, results for Go It Alone, Connected Communities, and Recent Trends are very similar and plot on top of each other. For the Cocheco River watershed, results for Go It Alone, Connected Communities, and land use in 2010 are very similar and plot on top of each other.

## 4 Discussion

### 4.1 Differences among land-use scenarios

Variations in the future land-use scenarios have little effect on the overall water balance and provisioning of streamflow. The dominant effect of land-use is on the temporal regulating service: partitioning streamflow between faster storm runoff and slower baseflow (Table 5 and Figure 4). The effects on these services are similar across the two climates and two watersheds. Increases in urban areas lead to more water moving quickly to the streams, which increases the magnitude of the annual maximum daily discharge. This effect reaches a maximum of approximately 10% for both the Cocheco and Charles River watersheds when comparing land use in 2010 with the Growing Global scenario.

The relative sensitivity of AMDF to impervious area is 2% for the Cocheco River watershed and 6% for the Charles River watershed (for both historic weather and a future climate). Thus, a large change in impervious area is required to generate a noticeable effect on the annual maximum daily flow (Table 8). These results are consistent with those of Hantush and Kalin (2006) who found a relative sensitivity of AMDF to developed area of 2% in Pennsylvania. Part of the reason for these limited sensitivities may be that high flows in New England and the northeast occur predominantly in March and April when evapotranspiration is low and the ground is saturated. Under such conditions, the regulating service associated with infiltration is reduced. Sensitivity of AMDF to precipitation is much greater: 40-60% for the Cocheco River watershed and over 80% for the Charles River. Even though the sensitivities are quite different, the effects on AMDF of plausible future changes in land use or climate in 2060 are comparable, with effects due to land-use change reaching 10% and effects attributable to climate change of approximately 5% for the Cocheco River and 17% for the Charles River (Table 8).

### 4.2 Implications of findings for policy and design

The results of this work indicate that the effects of climate and land use on runoff and high flows are additive (Table 8). The combination of a wetter future climate and increased urbanization has the potential to exacerbate high flows and flooding. While the results imply that it would take a major reworking of the landscape to mitigate the effects of climate change, they also indicate that rapid growth and development could present significant challenges for stormwater management and existing infrastructure. If population growth is modest, land-use decisions and development patterns have little effect on storm runoff and high flows (compare scenarios CC and GA in Table 5 and Figures 6 and 7). However, when the future is characterized by global socio-economic connectedness and increased population growth (Table 1), the results from the Yankee Cosmopolitan and Growing Global scenarios are substantively different (Table 5 and Figures 6 and 7). In this case, urban planning and choices regarding land use can have a large impact on regulating services and the potential for flooding. Planning for smart and sustainable growth while concomitantly investing in multi-functional landscapes and natural infrastructure could reduce flood damages. Additionally, with increased high flows, communities may need to increase the size of their water infrastructure and/or allow for short periods of inundation (Rosenzweig et al., 2018).

### 4.3 Limitations of approach

This study employs a hydrologic model to investigate the potential impacts of future land-use scenarios on streamflow. As such, the utility of the results depends upon the appropriateness of the mathematical representation of watershed characteristics and processes. SWAT is a well-established model, suitable for watershed applications, that has been and continues to be employed in a number of studies and investigations. Nonetheless, there are some inherent limitations of the model, and the results of this work should be interpreted within that context.

First, some of the model parameters (such as available water content, hydraulic conductivity, and surface runoff lag) are determined by calibrating the model to existing conditions. Using the model to represent future land use presumes that those parameters are unchanging across the scenarios. In most cases, we anticipate this to be true, as those parameters are functions of soil, topography, or other watershed characteristics that are generally unchanged as the land cover changes. Characteristics that do change with land use, such as the curve number and vegetation cover, are not calibrated but determined a priori. Second, the temporal resolution of this work is limited to the daily timescale. This precludes the representation of sub-daily dynamics of precipitation and streamflow. Therefore, instantaneous peak streamflows cannot be modeled, and this work is limited to daily discharge. Third, SWAT represents space in a semi-distributed way. While the model accounts for spatial variations among watershed characteristics, the HRU structure does not permit the representation of the spatial arrangement and connectedness of landscape elements. Therefore, feedbacks and interactions among different parts of the landscape cannot be represented explicitly. For example, increased runoff from one HRU cannot infiltrate in a different HRU. Such interactions can only be represented implicitly. Relatedly, storm runoff is represented with an approach that implicitly accounts for effects of soil, land cover, and land management through a single parameter. This is consistent with large-scale analyses and is not intended for small-scale green-infrastructure evaluation. Results from this work must be interpreted within the context of these modeling limitations.

### 4.4 Next steps

Our results reveal that potential changes to high flows are strongly connected to increases in urban land uses in New England. To more precisely elucidate the effects of such changes in land use and land cover, one could refine the representation of urban hydrology. Models such as the Storm Water Management Model (SWMM) and HydroCAD are better equipped to represent the natural and engineered features of an urban landscape, the sub-daily dynamics of the runoff response to storm events, and the elements of green infrastructure at the site and local scales. Such site-scale and sub-daily simulations of hydrological responses can further inform policy and practice, and these more detailed studies will necessarily be narrower in geographic scope. Continued engagement with stakeholders in the scenario-planning process can provide guidance to locations of interest along with the level of risk and types of landscape and infrastructure interventions that communities are willing to accept.

Finally, changes to nutrient and sediment loads are additional effects of changes to the landscape that may be of interest to stakeholders in New England. SWAT could be employed (for the Charles River, Cocheco River, or other watersheds) to investigate the effects of the landscape scenarios on the export of nitrogen, phosphorus, and sediment. The effects on water quality could be combined with our results on high flows to create a more complete picture of the effects of landscape futures on water-related services.

## 5 Conclusions

Application of a hydrologic model to stakeholder-developed scenarios can provide meaningful insight to the effects of plausible land-use changes on water-related ecosystem services. The combination of land use and climate change on storm runoff, high flows, and flooding are issues of concern, not only in New England but worldwide. Across the NELF scenarios, variations in land use had little effect on the overall water balance. Rather, the impact was on high flows and the partitioning of streamflow between storm runoff and baseflow. Those effects were correlated with the amount of impervious cover. For most of the scenarios (GA, CC, YC), the effects were muted and less than the effects due to climate change. For the Growing Global scenario, however, the effects were large and comparable to or greater than the effects of climate. These responses to land-cover change were similar across the Cocheco River and Charles River watersheds. Results from this work can help inform designs and decisions related to infrastructure resiliency and can complement other studies to provide a comprehensive assessment of ecosystem services across possible future landscapes.

## Acknowledgements

This research was supported, in part, by Highstead, the National Science Foundation Harvard Forest Long Term Ecological Research Program (Grant No. NSF-DEB 18-32210), and the Scenarios Society and Solutions Research Coordination Network (Grant No. NSF-DEB-13-38809). The authors would also like to thank Dr. James Dennedy-Frank for his helpful comments.

